# Control without cause: How covariate control biases our insights into brain architecture and pathology

**DOI:** 10.1101/2024.04.12.589152

**Authors:** Christoph Sperber, Laura Gallucci, Marcel Arnold, Roza M. Umarova

## Abstract

Inferential analysis of normal or pathological brain imaging data – as in brain mapping or the identification of neurological imaging markers – is often controlled for secondary variables. However, a rationale for covariate control is rarely given and formal criteria to identify appropriate covariates in such complex data are lacking. We investigated the impact and adequacy of covariate control in large-scale imaging data using the example of stroke lesion-deficit mapping. In 183 stroke patients, we evaluated control for age, sex, hypertension, or lesion volume when mapping real or simulated deficits. We found that the impact of covariate control varies and can be strong, but it does not necessarily improve the precision of results. Instead, it systematically shifts results towards the inversed associations between imaging features and the covariate. This effect of covariate control can bias results and, as shown in another experiment, can even create effects out of nothing. The widespread use of covariate control in the statistical analysis of clinical brain imaging data – and, likely, other biological high-dimensional data as well – may not generally improve statistical results, but it may just change them. Therefore, covariate control constitutes a problematic degree of freedom in the analysis of brain imaging data and may often not be justified at all.

## 1 Introduction

“Trust is good, control is better” could serve as a credo for many of the life sciences, including neuroscience: statistical control appears to be omnipresent, generally desirable and necessary. In imaging studies on human brain function or pathology, a plethora of variables is commonly controlled for as covariates, while only a minority of studies does not include any covariates.^1^ Easily accessible covariates such as age and sex are very common.^1^ Education level is a covariate in studies on cognitive and even non-cognitive target measures.^2–4^ The relationship between stroke lesion imaging and cognitive deficits or clinical outcomes is often controlled for lesion size.^5–8^ When mapping the neural correlates of different semantic error types in aphasia, taxonomic and thematic errors are controlled for each other.^9^ The association of small vessel disease markers and neurological health is investigated with 12 covariates including diabetes, smoking, and serum total cholesterol.^4^ The association of cortical thickness and the Big Five personality factors is controlled for intelligence, and each personality factor is controlled for the remaining four.^10^

Even only given this small number of examples, the question arises as to what criteria, aside from mere conventions of each field, should be used to select covariates. What covariates are included markedly varies even across similar studies, and brain mapping with between none and up to over a dozen covariates are publishable^1^. To our knowledge, a rationale for why a particular covariate is considered and why others are not is rarely stated. However, the inclusion of covariates can have different and strong effects.^1^ Hence, a clear, objective rationale for the selection of covariates seems necessary. Even more, control can have negative effects by overshadowing true associations or creating statistical artefacts. For example, a humorous illustration of the abuse of researchers’ degrees of freedom^11^ suggested that, after inclusion of a covariate, listening to a song biologically rejuvenates listeners. In stroke lesion-deficit mapping, control for lesion size or secondary deficits can create spurious associations.^12,13^

The literature on causal inference cannot only provide a theoretical background as to when and why to apply statistical control, but it also shows awareness of the potential dangers of ill-applied covariate control.^14–16^ When we want to infer causal effects from observational data, the strategy to identify a covariate requires making assumptions about the causal relationships between all variables. This strategy also identifies variables that should not be controlled for because they potentially mask true causal relationships or introduce spurious correlations. These causal assumptions are commonly visualised in causal graphs (for an accessible introduction, see^15,16^).

With the present manuscript’s focus on neuroscience, we should now realise the methodological abyss that has just opened up before us: in neurological or biological big data, statistical analyses often include thousands of variables. Let us look at the example of stroke lesion-deficit mapping, which investigates the relationship between the location of brain lesions and behavioural or cognitive deficits. The many imaging variables – like the lesion status of imaging voxels or brain regions – are not independent but are intertwined due to the underlying complex anatomy of the brain vasculature,^17^ where an occlusion within the hierarchy of the arterial structure *causes* pathology in many imaging features. Post-stroke deficits are *caused* by damage to certain brain structures and networks – but the severity of a deficit may vary across the location and extent of damage to relevant structures and may depend on varying interactions of multiple structures.^18^ Next, covariates can have a complex relationship with many variables. For example, consider sex: stroke outcome is linked to sex-specific lesion patterns,^19^ sex is associated with stroke outcomes and interacts with age,^20,21^ and animal models and theories on the impact of sex hormones suggest a *causal effect* of sex on stroke outcomes.^21–23^ Last, a *causal relationship* between sex and a clinical or cognitive target variable may exist through sociological, psychological or biological effects. In summary, it is difficult, if not impossible, to make reasonable assumptions about the causal relationships among all variables in brain mapping with tens to millions of variables. Hence, the requirement to evaluate the appropriateness of covariate control in brain mapping can hardly be fulfilled.

In the current study, we examined the impact and adequacy of common demographic or clinical covariates when mapping neurological and cognitive deficits. We focussed on the method of lesion-deficit mapping in stroke,^24^ as it is a popular method to infer functional brain architecture and as it can be objectively evaluated in silico.^25^ To gauge the effects of covariate control, we investigated i) how covariates affect real lesion-deficit mapping results, ii) how these covariates relate to lesion anatomy, iii) how covariates affect lesion-deficit mapping of in silico variables, i.e. simulated variables for which the neural correlates are perfectly known, and iv) the impact of covariates alone under the null hypothesis, i.e. when the mapped variables are not related to lesions of any brain area.

## 2 Methods

### 2.1 Patients and clinical assessment

We analysed data from a prospective study on post-stroke cognitive impairment^26^ (ClinicalTrials.gov Identifier: NCT05653141). This study recruited acute ischemic stroke patients admitted to the Stroke Centre of the University Hospital Bern. Detailed inclusion and exclusion criteria are reported elsewhere.^26^ In short, patients with a first-ever anterior circulation ischemic stroke confirmed by MRI and stroke onset ≤10 days ago were included. Patients with aphasia, which limits neuropsychological testing, were excluded; therefore the final sample contained more right-sided strokes. Clinical and demographic information was retrieved from the clinical documentation, including the diagnosis of arterial hypertension. We received written informed consent from all participants. The study was approved by the local ethics committee (Kantonale Ethikkommission Bern KEK 2020-02273).

A trained psychologist performed a multi-domain cognitive assessment during the hospital stay within 10 days after stroke onset. The present study focussed on the impact of covariates such as age or sex. Hence, we used raw performance scores and, contrary to common clinical practice, not scores that already account for such variables by age- or sex-specific norms. For the current study, we investigated three different neuropsychological measures: i) *Visuoconstructive ability* as included in the CERAD battery^27^. Patients were asked to copy four line drawings of increasing complexity and their performance was rated on an 11-point scale according to 2–4 pre-defined evaluation criteria for each shape. ii) Spatial *Selective attention* as included in the paper-and-pencil Bells Test^28^. Patients were asked to circle bell-shaped black icons placed among distractor items. The final score was the total number of omitted targets. iii) *Short-term memory* measured by digit span forward performance included in the WAIS-IV^29^. Patients were asked to repeat between 2 to 9 digits in the same order as read aloud by the examiner. The test was terminated after two false digit sequences in succession with the same length. The final measure was the maximum number of correctly repeated digits.

### 2.2 Brain imaging and lesion delineation

We identified lesioned tissue using diffusion-weighted MRI acquired 24h after stroke onset with MATLAB and SPM12 (https://www.fil.ion.ucl.ac.uk/spm/software/spm12/). The lesion area was delineated by a semiautomatic algorithm under supervision of a neurologist with >10 years of experience in lesion mapping (author R.M.U.) as done previously.^30^ Lesion masks were normalized to the Montreal Neurological Institute space at 2×2×2 mm³ resolution with normalisation parameters from the co-registered T1 scans. These procedures resulted in a binary mask for each patient indicating lesioned vs. intact voxels in a standardized coordinate system that allows spatial comparison of the lesion area between patients. Lesion volume was assessed as the volumetric size of each patient’s normalised binary mask.

### 2.3 Lesion-deficit mapping

We mapped lesion-deficit associations using well-established methods^24^ that statistically map the association between lesion status (lesioned vs. intact) and a deficit for each voxel in the brain. Analyses were done either with frequentist or Bayesian general linear models. *Frequentist statistical mapping* was performed in NiiStat (https://www.nitrc.org/projects/niistat) in MATLAB R2023a with control for multiple comparisons (i.e. control for α-error accumulation due to repeated testing in each voxel) by maximum statistic permutation, a family-wise error correction procedure^31^ with 4000 permutations and a one-sided α=0.05. Covariate control was implemented by the Freedman-Lane procedure,^32^ which allows for covariate control in a permutation design. *Bayesian lesion-deficit inference* (BLDI) was performed with the BLDI toolkit^13^ in R software 4.2.1 using Bayesian general linear models in the BayesFactor package v0.9.12^33^. Covariates were included as an additional independent variable. Resulting Bayes Factors indicate the ratio of evidence for the alternative hypothesis H1 (a lesion-deficit association exists, two-sided) against the null hypothesis H0 (a lesion-deficit association does not exist) and were interpreted according to common conventions^34^. All analyses were restricted to voxels that were damaged in at least 4 patients.

As default, we utilised frequentist statistical mapping, which is commonly considered a comparatively precise gold standard of mass-univariate voxel-wise mapping.^35^ We instead used BLDI whenever the focus of an experiment was the null hypothesis as Bayesian statistics, contrary to frequentist statistics, can provide evidence for the null hypothesis. We also used BLDI if real-world deficits yielded no significant results with the frequentist approach, as it can detect subtle effects^13^.

### 2.4 Study design and experiments

#### 2.4.1 Experiment 1 - Impact of covariate control on real-world lesion-deficit inference

We mapped the real-world post-stroke neuropsychological measures of visuoconstructive ability, selective attention, and short-term memory. We did so either without any covariate control, or while controlling for one of the following variables: age, sex, presence of hypertension, or lesion volume. We aimed to descriptively explore the impact of covariate control in such a setting. For each analysis, patients with missing neuropsychological measures were excluded (not more than 2 patients per variable).

#### 2.4.2 Experiment 2 - Associations of covariates and lesion status

Here, we focussed on the four covariates that were controlled for in the previous experiment – age, sex, hypertension, and lesion volume. We explored if these variables and the voxel-wise lesion status are associated. For each voxel in the brain, the Spearman rank correlation coefficient between lesion status (0 - intact; 1 - lesioned) and the variable of interest was computed.

#### 2.4.3 Experiment 3 - Impact of covariate control in a simulation with known ground truth

In this experiment, we aimed to evaluate covariate control in a perfectly transparent setting with a known ground truth which allows us to evaluate the precision of brain mapping. We followed a simulation strategy that has been used previously to evaluate the validity and precision of lesion-deficit inference.^25,35^ The basic concept is accessibly described in the Supplementary. Importantly, such simulation allows us to evaluate the impact of control of *real covariates* with *real imaging* data in a well-known system.

In the present experiment, we randomly placed a sphere with a 1cm radius in the tested brain space as a ground truth region. In the first condition, every patient whose lesion overlapped with this sphere was considered to suffer a deficit in the simulated variable. Hence lesion location, independent of covariates, was the only cause for the simulated deficit. Patients with the deficit received a random normal score of 0±1; patients without the deficit a score of 1.5±1. The effect size was optimised in piloting simulations to generate statistical maps with a size that neither extremely over-nor underestimates the size of the ground truth region. In the second condition, we simulated an impact of both damage to the ground truth region and the covariate. Again, scores of 0±1 were simulated for patients with damage to the ground truth region, and else 1.5±1. However, the mean value was also influenced by the relevant covariate. For sex or hypertension, the mean score was further decreased by 0.5 points, i.e., more pathological, for patients of male sex or with hypertension. For age, we computed the z-standardisation and subtracted z(age)*0.5 points from the score, i.e., higher age led to a more pathological performance. We did not include the covariate lesion volume in the second condition of experiment 3, as it would not conceptually fit the experimental design. The causal relationship between lesion volume and a deficit is complex, with strong associations between lesion volume and a deficit even without implicitly modelling such in a simulation^36^. For simplicity and interpretability, we always controlled only for one of the covariates. We repeated the simulation with 30 different ground truth spheres for each condition.

For the final analysis of the simulation experiment, we first compared several parameters between brain mapping with versus without covariate control: total number of significant voxels, positive predictive value (true positives/(true positives+false positives)), and number of false negatives. Variables were compared for each covariate separately with Wilcoxon-signed-rank tests with Bonferroni correction for multiple comparisons. In the second step, we investigated the influence of covariate control at the topographic level. This analysis focussed on the Z-statistic in the general linear model that is mapped across the brain. For each simulation, we computed a ΔZ map by subtraction of the map without covariate control from the map with covariate control. Averaging this map across all simulations shows us if and how the impact of covariate control is specific to certain regions. Further, with this map, we examined the relationship between the change in the mapped statistic after covariate control and the voxel-wise associations of covariates and lesion status obtained in experiment 2.

#### 2.4.4 Experiment 4 - Impact of covariate control under the null hypothesis

This experiment evaluated the impact of covariates under the null hypothesis, i.e., when associations between a behavioural measure and lesion location do not exist. We ensured that the null hypothesis was true by design using random data, which were generated based on the real neuropsychological data for visuoconstructive ability. For each simulation, we shuffled the original scores to create a measure with the distribution of a real neuropsychological measure, but for which any lesion-deficit association does not exist. We mapped lesion-deficit associations with BLDI and computed the extent of evidence for the null hypothesis H0 (i.e., voxels with Bayes factors ≤1/3^34^) and for the alternative hypothesis H1 (i.e., voxels with Bayes factors ≥3). We performed 30 simulations of a behavioural measure and mapped each measure either without a covariate or one of the four potential covariates.

### 2.5 Data availability

Statistical maps and scripts are available at https://data.mendeley.com/datasets/gvn3k8cj9r/1. Original clinical data are not publicly available, but qualified researchers may request access to anonymised data. Proposals need to be approved by the local ethics commission.

## 3 Results

We analysed data from 183 acute stroke patients. Information on sample demographics and study variables is shown in Supplementary Table 1. The correlation of neuropsychological measures and potential covariates varied (Table 1). Visuoconstructive ability correlated with all potential covariates, selective attention with age and lesion volume, and short-term memory only with lesion volume. The correlation strength varied from weak to strong (Spearman’s ρ=0.55). We also found correlations between some of the covariates: sex and age correlated weakly, with men being younger, and sex and hypertension correlated strongly, with men suffering more often from hypertension. We performed statistical mapping for 31206 voxels that were lesioned in at least 4 patients (see Supplementary Figure 1).

**Table 1:** Correlations between neuropsychological measures and deficits. Spearman’s rank correlation between study variables. Binary variables were coded by 0/1. Asterikses indicate significance at * p<0.05; ** p<0.01; *** p<0.001.

|  | Age | Sex (1=male) | Hypertension<br>(1=present) | Lesion<br>Volume |
| --- | --- | --- | --- | --- |
| Visuoconstructive ability | -0.307*** | 0.182* | 0.152* | 0.465*** |
| Selective attention | -0.181* | 0.136 | 0.051 | 0.551*** |
| Short-term memory | -0.042 | 0.058 | 0.087 | 0.181* |
| Age | 1 | -0.216** | -0.067 | -0.095 |
| Sex |  | 1 | 0.523*** | -0.109 |
| Hypertension |  |  | 1 | -0.022 |
| Lesion Volume |  |  |  | 1 |

### 3.1 Experiment 1 - Impact of covariate control on real-world lesion-deficit inference

The impact of covariate control on real-world deficits varied and occasionally created massive changes in statistical results (Figure 1). For *visuoconstructive ability*, the impact of control for age, sex, or hypertension was mostly quantitative. The clusters of voxels implicated without control were also present with covariate control, but, depending on the covariate and localisation, were enlarged or reduced in size. Occasionally, this change in the extent of clusters meant that some brain regions were additionally or not anymore implicated. The results after control for lesion volume stood out with rather small clusters that were entirely different from the uncontrolled results. For *selective attention*, control for age, sex, or hypertension modified the extent of most clusters and even removed some small clusters entirely. With control for lesion size, the analysis found almost no results anymore, except for a tiny, non-interpretable cluster of voxels in the white matter. Notably, after lesion size control, the frequentist mapping of both visuoconstructive ability and selective attention also found significant results that only appeared when testing the inverse association, i.e., the hypothesis that a lesion is linked to better and less pathological performance.

**Figure 1:**
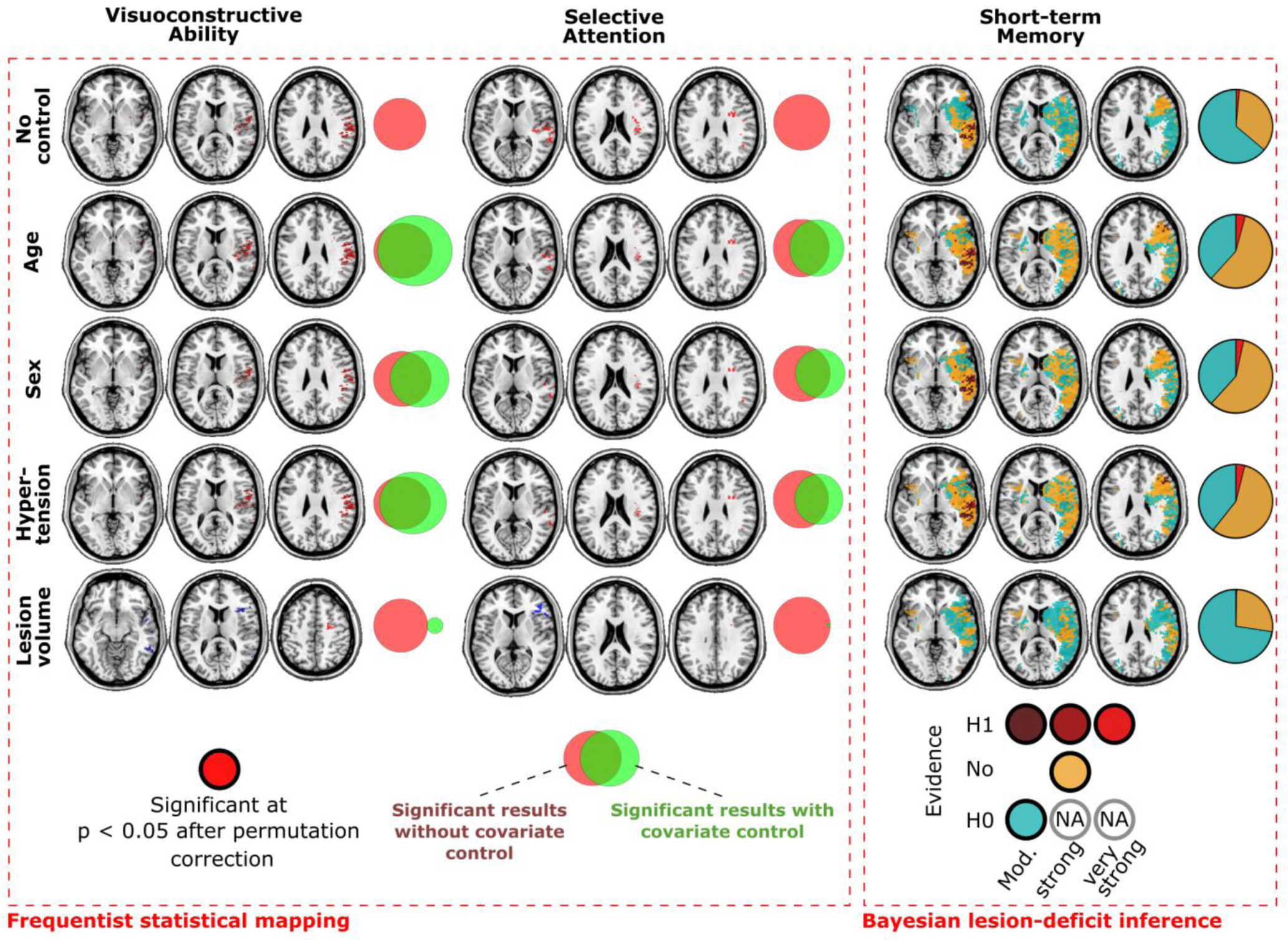
Results of Experiment 1. Lesion-deficit mapping of real-world post-stroke neuropsychological measures. For short-term memory, no statistically significant results were found with frequentist mapping under any condition and results of Bayesian mapping are shown instead. For frequentist analyses (left and middle column), Venn diagrams show the spatial overlap of the map without covariate control and each map with control; for Bayesian analyses, Venn diagrams show the proportion of tested voxels within each evidence category. For analyses with the covariate lesion volume, frequentist analyses found areas with inversed lesion-deficit associations (i.e. where lesions were associated with better performance), which are shown in blue.

For *short-term memory*, frequentist analysis generated null results under all conditions. We hence re-analysed the measure with Bayesian lesion-deficit inference (BLDI). The uncontrolled analysis found evidence for the null hypothesis in many brain regions, while evidence for the alternative hypothesis, i.e. lesion-deficit associations, was limited to inferior and middle temporal regions. With control for age, sex, or hypertension, results seemed to generally shift towards the alternative hypothesis. Evidence for the null hypothesis became scarcer and some new clusters with evidence for lesion-deficit associations emerged, for example in frontal areas and the white matter. Control for lesion volume appeared to have the opposite effect – larger areas with evidence for the null hypothesis were found while evidence for the alternative hypothesis decreased markedly.

### 3.2 Experiment 2 - Associations of covariates and lesion imaging features

For all variables that we controlled for in the previous experiment, we observed at least weak correlations (Spearman’s |ρ|≥0.1) with the voxel-wise lesion status in both the negative and positive direction (Figure 2). The lowest peak correlations were found for sex (|ρ|=0.23) and the highest for lesion volume (ρ=0.52). For age, sex, and hypertension, several clusters of voxels with at least weak correlations were found scattered across the brain. For lesion volume, correlations were higher, and the lesion status of almost all voxels in the brain correlated with lesion volume. In summary, the variables we included as covariates in Experiment 1 are associated with certain lesion locations, whether by chance or true associations between lesion anatomy and the variables.

**Figure 2:**
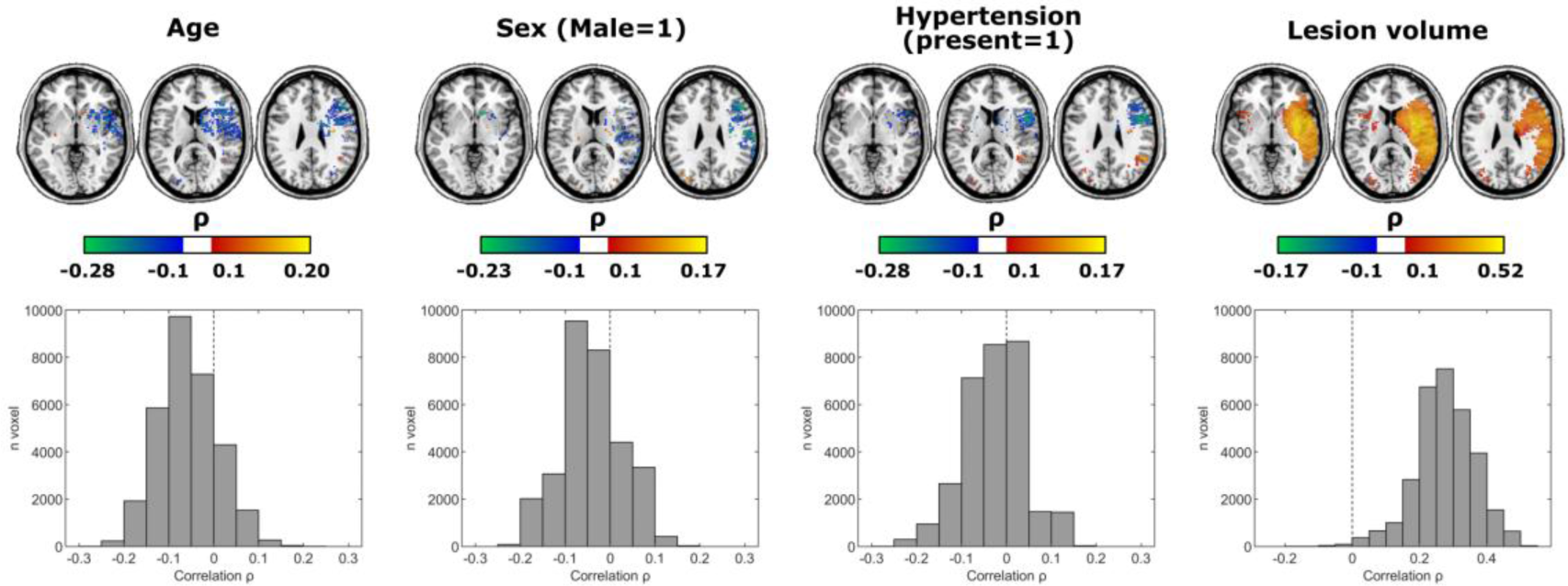
Results of Experiment 2. Voxel-wise correlation between lesion status and the variables controlled for in other experiments. Covariates were found to be associated with the lesion status of several brain regions. The upper panels show all voxels with a Spearman’s rank correlation of at least |ρ|≥0.1. Colour scales represent the range from |ρ|=0.1 to the maximum |ρ| scores. The lower panels show the distribution of ρ values. The voxel-wise lesion status was coded as intact (0) or lesioned (1), sex as female (0) or male (1), and hypertension as absent (0) or present (1).

### 3.3 Experiment 3 - Impact of covariate control in a simulation with known ground truth

In the first condition of the simulation, only a direct causal impact of lesion location, but not the covariate was simulated. For the covariates age, sex, and hypertension, covariate control had no significant effect on any of the parameters – number of significant voxels, positive predictive value, and number of false negatives. With control for lesion volume, the number of significant voxels decreased strongly (Example in Figure 3A). The number of false negatives decreased accordingly, but the positive predictive value remained unchanged. For detailed statistics, see Supplementary Table 2. In line with the absence of effects, the maps of the average statistical changes through covariate control (Figure 3B) were very small and close to zero, except for lesion volume control. The average statistical changes with lesion volume control correlated strongly with the map of lesion - lesion volume associations generated in experiment 2 (ρ(31206)=-0.65; p<0.0001), meaning that the effect of lesion volume control was region-specific and depended on the focal relationship between lesion status and lesion size. In the second condition of the simulation, both a direct causal impact of lesion location and the covariate were simulated. Here, the total number of significant voxels increased when any covariate was controlled for. The number of false negatives minimally, but significantly increased, but the positive predictive value was unchanged. For detailed statistics, see Supplementary Table 3. The maps of the average statistical changes through covariate control (Figure 4C) now included much larger values. For the covariate age, the average change in statistical maps ranged from ΔZ=-1.06 to 1.65, which correlated almost perfectly (ρ(31206)=- 0.95; p<0.0001) with the association map generated in experiment 2. For sex, the average change in statistical maps ranged from ΔZ=-0.49 to 0.76, likewise with an almost perfect correlation with the map from experiment 2 (ρ(31206)=-0.99; p<0.0001). For hypertension, the average change in statistical maps ranged from ΔZ=-0.46 to 0.92 and again correlated almost perfectly (ρ(31206)=-0.99; p<0.0001). To put these Z scores into reference, the permutation-based threshold |Z| score to consider a statistical result significant in the uncontrolled analyses without any impact of the covariate was Z=4.15±0.14. In summary, i) covariate control changed statistical results, but not systematically for the better, and ii) lesion-covariate associations as shown in experiment 2 (Figure 2) seemed to almost perfectly describe the average effect of covariate control in lesion-deficit mapping. In brain regions for which a covariate was anticorrelated with the lesion status, covariate control shifted results towards assuming a lesion-deficit association. Vice versa, in regions for which the covariate was correlated with the lesion status, covariate control shifted results towards the null hypothesis.

**Figure 3:**
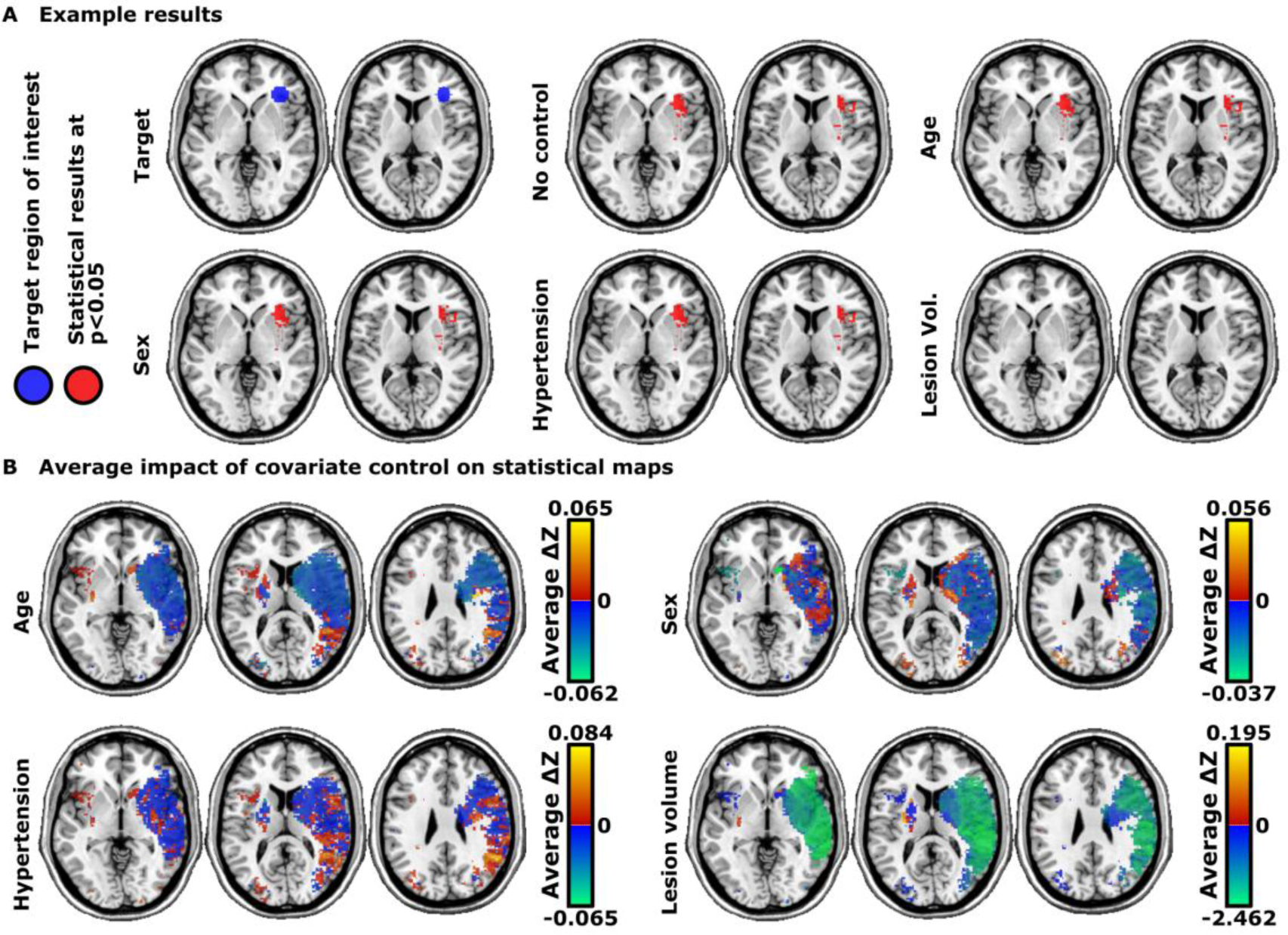
Results of Experiment 3 – Condition with simulated sole impact of lesion status. Covariate control for lesion volume had a strong impact, while the impact of other covariates was negligible. **(A)** Example results for a single simulation run.. **(B)** The average change in the voxel-wise Z-statistic after covariate control across 30 simulation runs. The indicated scaling of the values should be paid attention to; values were very small and indicated only a minimal impact of covariate control in this condition. The values were coded so that warm colours (red-yellow) show areas for which statistics are pushed towards suggesting a lesion-deficit association. To put these Z scores into reference, the average threshold to consider a statistical result significant in the uncontrolled analyses was |Z|=4.15±0.14.

**Figure 4:**
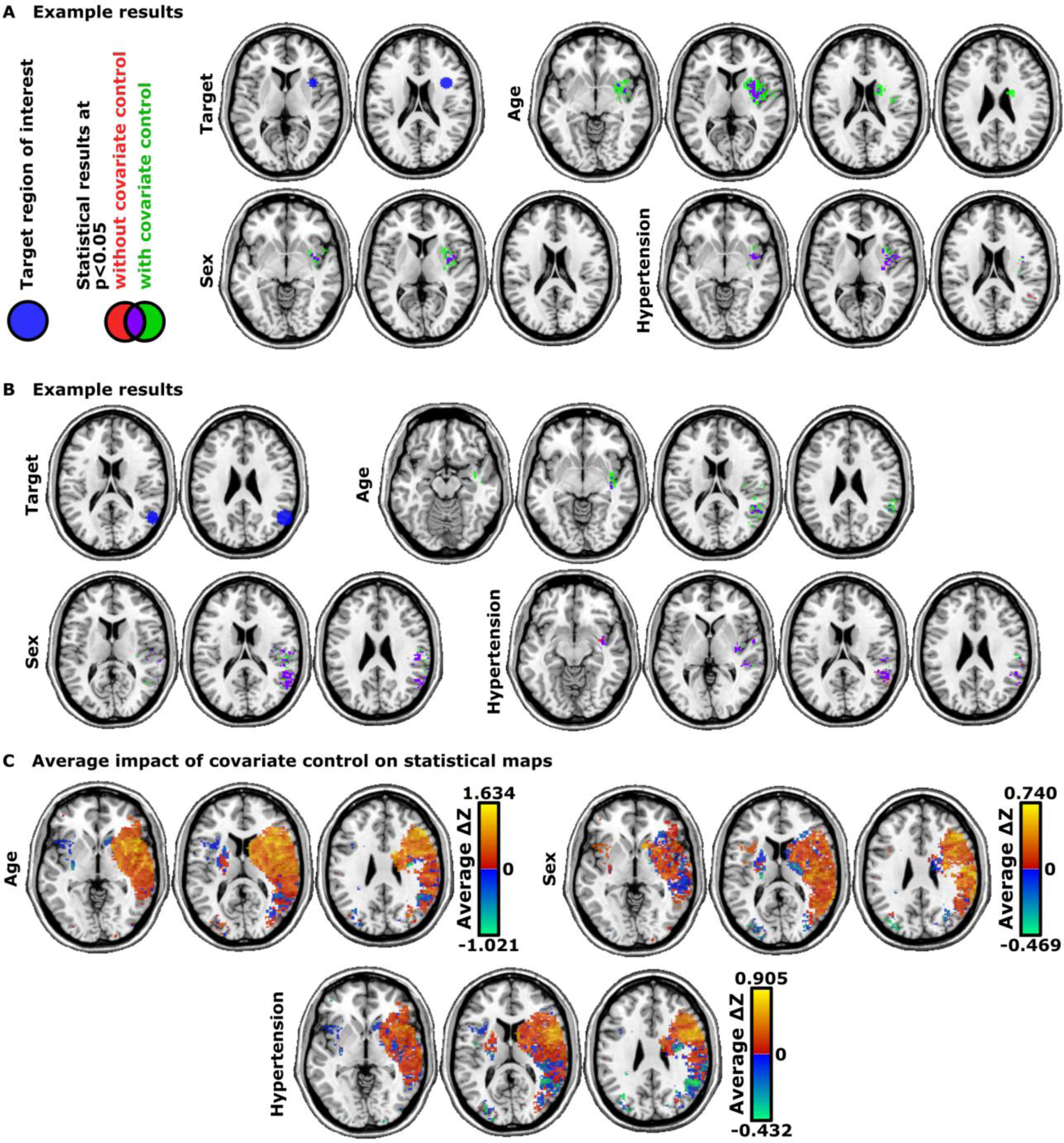
Results of Experiment 3 – Condition with simulated impact of both lesion status and covariate. Covariate control had a strong impact specific to certain regions. **(A+B)** Example results for a single simulation run. **(C)** The average change in the voxel-wise Z-statistic after covariate control across 30 simulation runs. The indicated scaling of the values should be paid attention to. The values were coded so that warm colours (red-yellow) show areas for which statistics are pushed towards suggesting a lesion-deficit association.

### 3.4 Experiment 4 - Impact of covariate control under the null hypothesis

In data for which the null hypothesis is true by definition, we mapped lesion-deficit associations by BLDI. Under this condition, Bayes factors should be as small as possible and smaller than 1/3, i.e., should provide at least moderate evidence in favour of the null hypothesis H0 over the alternative hypothesis H1 (following existing conventions^34^).

Accordingly, the majority of Bayes factors were smaller than 1/3 across all simulations and independent of any covariate. However, the inclusion of a covariate generally increased Bayes factors, meaning that they shifted evidence from being in favour of H0 towards being in favour of H1 (Figure 5). The number of voxels with at least moderate evidence for H0 decreased under any covariate control (Bonferroni-corrected Wilcoxon-signed-rank tests, all p<0.0001). The number of voxels for which Bayes factors incorrectly indicated at least moderate evidence in favour of H1 (Bayes factors > 3) increased (all corrected p<0.0045). Notably, lesion volume stood out among all covariates with the nominally highest impact

**Figure 5:**
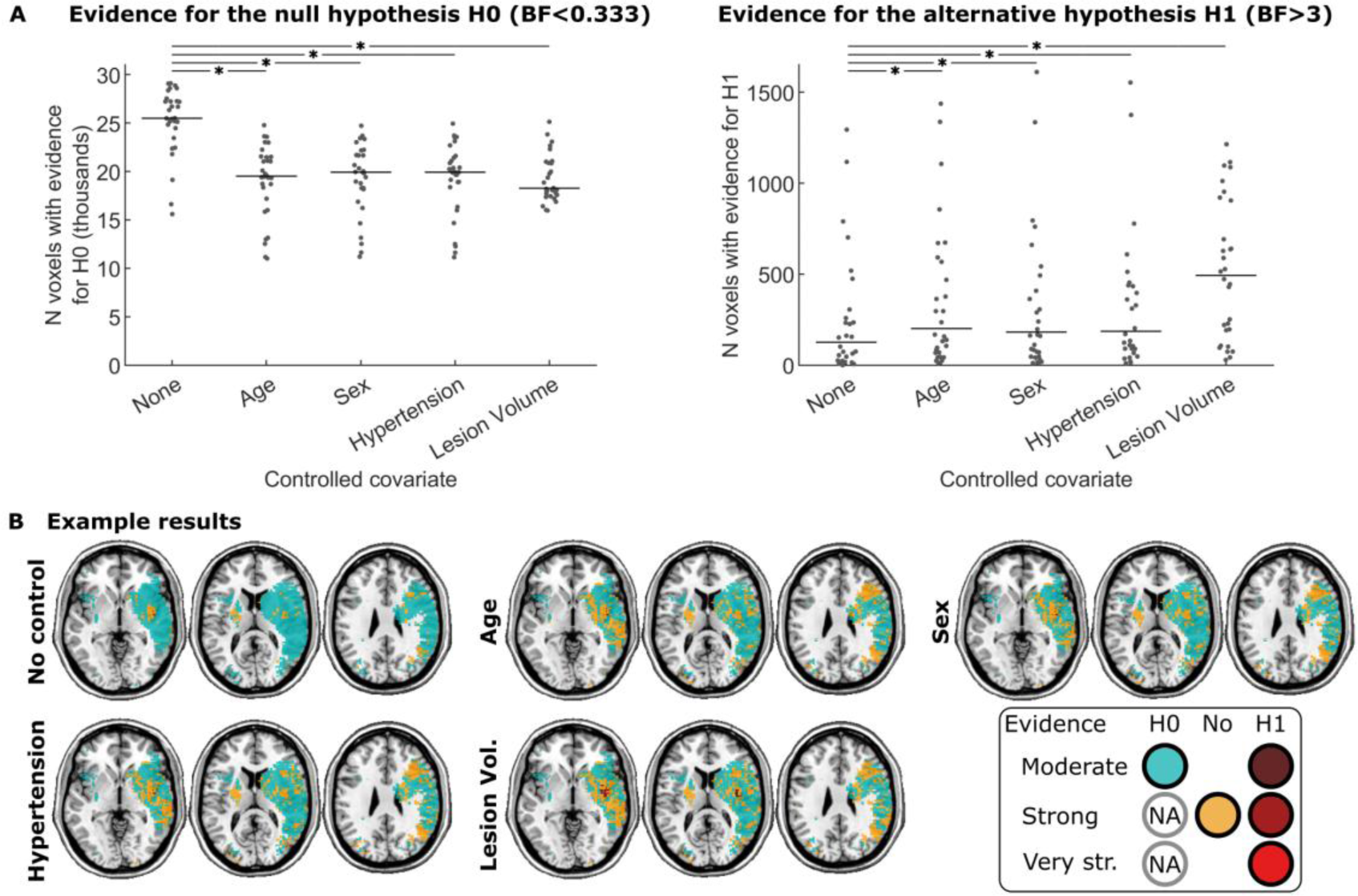
Results of Experiment 4. Covariate control falsely decreased the number of tests indicating evidence in favour of the null hypothesis. **(A)** Results of the simulation experiment under the null hypothesis. Each dot stands for one simulation, i.e. the statistical mapping of one simulated score. Bars show the median. Asterisks indicate a significant difference after Bonferroni correction. **(B)** Example results for a single simulation run.

## 4 Discussion

In multiple experiments, we evaluated the impact and adequacy of covariate control when mapping diseased brains in lesion-deficit mapping. In Experiment 1, we found that the effects of control for age, sex, presence of hypertension, and stroke lesion volume varied by covariate and poststroke deficit. While in some cases the effects were only minor, in other cases, they were more substantial and suggested partially different neural correlates. This leads us to the obvious question of which result – if any – corresponds to the truth. Without a precise rule for the selection of appropriate covariates, we face the serious epistemological problem that numerous contradictory results can exist. In Experiment 2, we have shown that the covariates are associated with the imaging data of brain pathology. Hence, in real data, associations may indeed be as complex as indicated in the introduction. Brain imaging variables and secondary variables that we control should not generally be seen as two factors that independently impact the genesis of behaviour or pathology, but as closely and complexly intertwined.

We have shown the consequences of this complexity in the next experiments. In Experiment 3, we evaluated the impact of covariate control in a well-controlled artificial setting. Covariate control had little to no impact when our simulations did not include a direct causal relationship of the covariate on the target variable. With such a causal relationship included, covariate control had an impact. Of note, we believe that such a direct causal effect likely represents the most common idea that researchers have about the impact of a covariate variable on behaviour or pathology. And, arguably, such effect alone – a simple linear effect independent of any imaging feature and without any interactions – should provide a relatively simple benchmark of what covariate control may accomplish. However, even though covariate control affected statistical results, we found no systematic improvement. Instead, what we found to be systematic was that covariate control changed statistical results following the inversed associations of covariates and lesion imaging features uncovered in Experiment 2. In other words: covariate control impacted results, but instead of improving them, it rather invertedly imposed the covariate’s association with the lesion anatomy onto statistical results. A researcher who, for example, controls stroke lesion-deficit mapping for the variable sex might do so to “control for the effect of sex”. This does not mean that the analysis better captures the neural correlates of a deficit. Instead, given the associations we found between sex and lesion status in

Experiment 2, it means that we effectively amplify statistical signal in frontal and temporal areas and attenuate signal in parietal areas. In Experiment 4, we found that covariate control can work against a confirmed null effect. In other words, in the absence of a true association between imaging features and a target variable, covariate control can erroneously shift results towards the alternative hypothesis and introduce spurious results. Hence, at least in the context of Bayesian statistics, the selection of inappropriate covariates can thus create statistical effects out of nothing.

To our knowledge, covariate control most often does not require justification in neuroimaging studies. On the contrary, we have the feeling that rather the omission of a statistical control that follows a convention of the respective field requires justification towards colleagues and reviewers. Considering the incentives in academia, this imbalance poses extreme dangers for scientific progress: if new, surprising results can achieve the greatest scientific prestige, arbitrary, unjustified covariate control provides an ultimate degree of freedom to be abused. While the general issue has already been discussed for many years,^11^ it is far more complex and magnified in large-scale data such as clinical brain imaging. Not only are there many independent variables (e.g., voxel-wise imaging features), but clinical routine diagnostics and documentation provide a large collection of potential covariates.

Among the four potential covariates that we evaluated in the current study, lesion volume stood out. The distinctive feature of this variable could be the high correlation with the imaging data. Experiment 2 found up to strong correlations between lesion volume and voxel-wise lesion status for almost the entire brain. Another feature could be the close and not clearly comprehensible coupling of the two variables: both lesion volume and lesion location are inseparably intertwined and it appears impossible to attribute the direct cause of a deficit to lesion volume alone.^36^ Covariates for which a causal relationship with other variables cannot be clearly defined are often encountered in neuroscience. We probably cannot clearly define a causal relationship between one personality trait and another. Then what is the neural correlate of a personality trait controlled for several other personality traits?^10^ What is a subscore in a stroke severity assessment battery controlled for the total test score?^37^ What is focal cortical thickness controlled for average brain-wide cortical thickness?^38^ Even if such covariates – at least situationally – improve statistical analysis, we still face the issue that causal relationships become even more difficult to grasp than they already are in large-scale neurobiological data. And whatever covariance control does will only become more complex and incomprehensible.

What variables should we control for in inferential analyses of biological/neurological large-scale data? Our current work is unable to provide any rule to answer this question and we doubt that any simple rule can be created at all. The sometimes utilized strategy to control all variables that are associated with the dependent variable appears to be invalid. The flaw in this approach is known in the literature on causal inference^14,16^ and, accordingly, we found that control for variables that are associated with a target variable – as by design in the second version of Experiment 3 – did not improve results. If anything, our data raises new, urgent questions. Even if covariate control could be appropriate, it might be difficult to select among strongly correlating potential covariates, such as age and hypertension in the current study. In this case, both variables could also be controlled, which leads us to the main limitation of the current study: we aimed to keep our simulations and analyses as simple as possible. We limited simulations to simple effects and only ever controlled for one covariate at once. And yet the complexity of the data was sufficient to produce undesirable changes in the results. But what if we step up the complexity and control for a dozen covariates, as done in actual studies?^1^ The issues identified in our study will probably not only persist in more complex situations but be severely amplified. Likewise, our simulations only looked at relatively simple causal relationships between a covariate and a target variable. Our study does not allow any strong conclusions to be drawn on what covariate control does in the face of complex interactions between covariates and imaging variables. Here too, we assume that the problems mentioned will only be exacerbated.

In conclusion, the widespread use of covariate control in the statistical analysis of clinical brain imaging data – and, likely, other biological high-dimensional data as well – may not generally improve statistical results, but it may just change them. Therefore, covariate control constitutes a problematic degree of freedom in the analysis of brain imaging data. The current modus operandi of selecting statistical covariates by correlation with the target variable, according to conventions, or plainly by availability, needs revision. However, given the high complexity of neurobiological data, the creation of objective rules to identify appropriate covariates might be difficult up to impossible. In any case, covariate control should be limited, and uncontrolled statistical results should also be considered in the interpretation of a study.

## Acknowledgements

This work is funded by the Synapsis Foundation (Grant number 2019-PI05) and the Heidi Seiler Foundation.

## Abbreviations

BLDI: Bayesian Lesion-deficit inference

## References

1. Hyatt, C. S. et al. The quandary of covarying: A brief review and empirical examination of covariate use in structural neuroimaging studies on psychological variables. Neuroimage 205, 116225 (2020).

2. Esteves, M. et al. Structural laterality is associated with cognitive and mood outcomes: An assessment of 105 healthy aged volunteers. Neuroimage 153, 86–96 (2017).

3. Evans, T. E. et al. Subregional volumes of the hippocampus in relation to cognitive function and risk of dementia. Neuroimage 178, 129–135 (2018).

4. Yilmaz, P., Ikram, M. K., Niessen, W. J., Ikram, M. A. & Vernooij, M. W. Practical small vessel disease score relates to stroke, dementia, and death: The Rotterdam study. Stroke 49, 2857–2865 (2018).

5. Mirman, D. et al. Neural organization of spoken language revealed by lesion-symptom mapping. Nat. Commun. 6, 6762 (2015).

6. Wu, O. et al. Role of Acute Lesion Topography in Initial Ischemic Stroke Severity and Long-Term Functional Outcomes. Stroke 46, 2438–2444 (2015).

7. DeMarco, A. T. & Turkeltaub, P. E. A multivariate lesion symptom mapping toolbox and examination of lesion-volume biases and correction methods in lesion-symptom mapping. Hum. Brain Mapp. 39, 4169–4182 (2018).

8. Klingbeil, J., Wawrzyniak, M., Stockert, A., Karnath, H. O. & Saur, D. Hippocampal diaschisis contributes to anosognosia for hemiplegia: Evidence from lesion network-symptom-mapping. Neuroimage 208, (2020).

9. Schwartz, M. F. et al. Neuroanatomical dissociation for taxonomic and thematic knowledge in the human brain. Proc. Natl. Acad. Sci. U. S. A. 108, 8520–8524 (2011).

10. Lewis, G. J. et al. Widespread associations between trait conscientiousness and thickness of brain cortical regions. Neuroimage 176, 22–28 (2018).

11. Simmons, J. P., Nelson, L. D. & Simonsohn, U. False-positive psychology: Undisclosed flexibility in data collection and analysis allows presenting anything as significant. Psychol. Sci. 22, 1359–1366 (2011).

12. Sperber, C., Nolingberg, C. & Karnath, H. O. Post-stroke cognitive deficits rarely come alone: Handling co-morbidity in lesion-behaviour mapping. Hum. Brain Mapp. 41, 1387–1399 (2020).

13. Sperber, C., Gallucci, L., Smaczny, S. & Umarova, R. Bayesian lesion-deficit inference with Bayes factor mapping: Key advantages, limitations, and a toolbox. Neuroimage 271, 120008 (2023).

14. Pearl, J., Glymour, M., & Jewell, N. P. Causal inference in statistics (1st ed.). (Wiley, 2016).

15. Rohrer, J. M. Thinking Clearly About Correlations and Causation: Graphical Causal Models for Observational Data. Adv. Methods Pract. Psychol. Sci. 1, 27–42 (2018).

16. Wysocki, A. C., Lawson, K. M. & Rhemtulla, M. Statistical Control Requires Causal Justification. Adv. Methods Pract. Psychol. Sci. 5, 251524592210958 (2022).

17. Zhao, L. et al. Strategic infarct location for post-stroke cognitive impairment: A multivariate lesion-symptom mapping study. J. Cereb. Blood Flow Metab. 38, 1299–1311 (2018).

18. Toba, M. N. et al. Revisiting ‘brain modes’ in a new computational era: Approaches for the characterization of brain-behavioural associations. Brain 143, 1088–1098 (2020).

19. Bonkhoff, A. K. et al. Sex-specific lesion pattern of functional outcomes after stroke. Brain Commun. 4, 1–12 (2022).

20. Reeves, M. J. et al. Sex differences in stroke: epidemiology, clinical presentation, medical care, and outcomes. Lancet Neurol. 7, 915–926 (2008).

21. Roy-O’Reilly, M. & McCullough, L. D. Age and sex are critical factors in ischemic stroke pathology. Endocrinology 159, 3120–3131 (2018).

22. Lisabeth, L. & Bushnell, C. Stroke risk in women: The role of menopause and hormone therapy. Lancet Neurol. 11, 82–91 (2012).

23. Herson, P. S. et al. Experimental pediatric arterial ischemic stroke model reveals sex-specific estrogen signaling. Stroke 44, 759–763 (2013).

24. Rorden, C. & Karnath, H. O. Using human brain lesions to infer function: A relic from a past era in the fMRI age? Nat. Rev. Neurosci. 5, 812–819 (2004).

25. Mah, Y. H., Husain, M., Rees, G. & Nachev, P. Human brain lesion-deficit inference remapped. Brain 137, 2522–2531 (2014).

26. Gallucci, L. et al. Post-stroke cognitive impairment remains highly prevalent and disabling despite state-of-the-art stroke treatment. International Journal of Stroke. In press.

27. Morris, J. C. et al. The consortium to establish a registry for alzheimer’s disease (CERAD). Part I. Clinical and neuropsychological assessment of alzheimer’s disease. Neurology 39, 1159–1165 (1989).

28. Gauthier, L., Dehau Lab Th-Alajouanine, F. T. & Yves Joanette, M. The Bells Test: A Qua ntitative a nd Qua lita ti ve Test For Visual Neglect. Int. J. Clin. Neuropsychol. **XI**, 49–54 (1989).

29. Wechsler, D. Wechsler Adult Intelligence Scale--Fourth Edition (WAIS-IV). (APA PsycTests, 2008).

30. Umarova, R. M. et al. Cognitive reserve impacts on disability and cognitive deficits in acute stroke. J. Neurol. 266, 2495–2504 (2019).

31. Nichols, T. E. & Holmes, A. P. Nonparametric permutation tests for functional neuroimaging: A primer with examples. Hum. Brain Mapp. 15, 1–25 (2002).

32. Winkler, A. M., Ridgway, G. R., Webster, M. A., Smith, S. M. & Nichols, T. E. Permutation inference for the general linear model. Neuroimage 92, 381– 397 (2014).

33. Morey, R.D., Rouder, J.N., Jamil, T., Urbanek, S., Forner, K., & Ly, A. Bayes Factor (Version 0.9.12-4.4) (2018). Retrieved from https://CRAN.R-project.org/package=BayesFactor

34. Wagenmakers, E. J. et al. Bayesian inference for psychology. Part II: Example applications with JASP. Psychon. Bull. Rev. 25, 58–76 (2018).

35. Pustina, D., Avants, B., Faseyitan, O. K., Medaglia, J. D. & Coslett, H. B. Improved accuracy of lesion to symptom mapping with multivariate sparse canonical correlations. Neuropsychologia 115, 154–166 (2018).

36. Sperber, C. The strange role of brain lesion size in cognitive neuropsychology. Cortex 146, 216–226 (2022).

37. Cheng, B. et al. Mapping the deficit dimension structure of the National Institutes of Health Stroke Scale. eBioMedicine 87, 104425 (2023).

38. Golchert, J. et al. Individual variation in intentionality in the mind-wandering state is reflected in the integration of the default-mode, fronto-parietal, and limbic networks. Neuroimage 146, 226–235 (2017).

