## Supplementary for "Control without cause: How covariate control biases our insights into brain architecture and pathology"

### Basic concept of the simulation approach in Experiment 3

The basic concept of the simulation design used in Experiment 3 is as follows: firstly, lesion masks are generated in a real data sample of stroke patients. These indicate the lesioned area in each patient. Secondly, the researcher defines a ground truth that explains how a fictional, simulated deficit is caused by neural damage. For example, one could decide that any lesion to the supramarginal gyrus causes a deficit. Such a rule is applied to simulate a score for each patient based on each lesion map. Thirdly, the neural correlates of the simulated score are mapped onto the brain with a lesion-deficit inference method. Lastly, the precision of the statistical map is evaluated. In the above-mentioned example, valid lesion-deficit inference should identify the supramarginal gyrus as the neural correlate of the deficit, and any systematic deviation uncovers the limitations of the inference method.

**Supplementary Table 1: Descriptive statistics**

IQR – interquartile range; SD – standard deviation.

| | All participants, N = 183 |
|---|---|
| Age, years mean (SD; range) | 65.8 (14.8; 26-98) |
| Sex Female/Male, % | 42.1/57.9 |
| Lesion size in $cm^3$, median(IQR; range) | 2.7 (0.8; 13.4; 0.1-251.5) |
| Hypertension, % | 67.8 |
| Visuoconstructive ability score, mean (SD; range) | 9.1 (1.8; 2-11) |
| Selective attention, number of omissions, mean (SD; range) | 4.7 (6.6; 0-33) |
| Short-term memory, maximal correctly repeated digit span, mean (SD; range) | 6.0 (1.7; 2-11) |

**Supplementary Table 2: Statistics Experiment 3 – Condition 1**

Detailed results of statistical analyses in experiment 3 for the first condition where only an impact of lesion location was simulated. Statistical comparison was performed by Wilcoxon Signed-rank tests. Reported p-values were corrected by the Bonferroni algorithm to account for multiple comparisons across 4 covariates. Asterisks indicate significance at * p<0.05; ** p<0.01; *** p<0.001. IQR – interquartile range.

| Covariate | Median(IQR) | Statistical comparison to no control condition |
|---|---|---|
| Total number of significant voxels | | |
| No control | 548 [128; 1697] | - |
| Age | 540 [149;1613] | *Z*=-0.75, *p*=1 |
| Sex | 540 [112; 1637] | *Z*=-1.86, *p*=0.25 |
| Hypertension | 653 [123; 1598] | *Z*=-0.48, *p*=1 |
| Lesion Volume | 11 [4; 98] | *Z*=-4.70, *p*< **0.0001*** ** |
| Positive predictive value (=Sensitivity) | | |
| No control | 0.06 [0.02; 0.22] | - |
| Age | 0.06 [0.02; 0.22] | *Z*=0.95, *p*=1 |
| Sex | 0.06 [0.02; 0.21] | *Z*=0.50, *p*=1 |
| Hypertension | 0.06 [0.02; 0.21] | *Z*=1.04, *p*=1 |
| Lesion Volume | 0.04 [0.00; 0.17] | *Z*=1.43, *p*=0.61 |
| Number of false negatives | | |
| No control | 307 [218; 396] | - |
| Age | 295 [220; 395] | *Z*=-0.42, *p*=1 |
| Sex | 325 [215; 398] | *Z*=-1.17, *p*=0.97 |
| Hypertension | 320 [212; 394] | *Z*=0.02, *p*=1 |
| Lesion Volume | 375 [243; 481] | *Z*=-4.54, *p*< **0.0001*** ** |

**Supplementary Table 3: Statistics Experiment 3 – Condition 2**

Detailed results of statistical analyses in experiment 3 for the second condition where both an impact of lesion location and the covariate were simulated. Statistical comparison was performed by Wilcoxon Signed-rank tests. Reported p-values were corrected by the Bonferroni algorithm to account for multiple comparisons across 3 covariates. Asterisks indicate significance at * p<0.05; ** p<0.01; *** p<0.001.

| Covariate | Baseline without covariate control Median(IQR) | With covariate control Median(IQR) | Statistical comparison |
|---|---|---|---|
| Total number of significant voxels | | | |
| Age | 55 [11;298] | 484 [27; 1818] | *Z*=-4.62, *p*< **0.0001*** ** |
| Sex | 223 [14; 1473] | 411 [45; 1850] | *Z*=-4.62, *p*< **0.0001*** ** |
| Hypertension | 198 [56; 506] | 307 [75; 724] | *Z*=-3.94, *p*< **0.0001*** ** |
| Positive predictive value (=Sensitivity) | | | |
| Age | 0.03 [0.01; 0.08] | 0.04 [0.02; 0.12] | *Z*=-0.86, *p*=1 |
| Sex | 0.05 [0.01; 0.12] | 0.07 [0.03; 0.12] | *Z*=-0.26, *p*=1 |
| Hypertension | 0.04 [0.00; 0.09] | 0.03 [0.00; 0.11] | *Z*=0.99, *p*=0.97 |
| Number of false negatives | | | |
| Age | 366 [224; 491] | 351 [191; 427] | *Z*=3.61, *p*= **0.0009*** ** |
| Sex | 324 [190; 439] | 304 [201; 426] | *Z*=3.49, *p*= **0.0014*** ** |
| Hypertension | 348 [230; 445] | 322 [214; 444] | *Z*=2.69, *p*= **0.0215*** |

**Supplementary Figure 1: Lesion overlap**

**(A)** Overlap topography of the total sample of 183 patients. **(B)** The same topography thresholded at n≥4, i.e. the area that was included in brain mapping analyses.

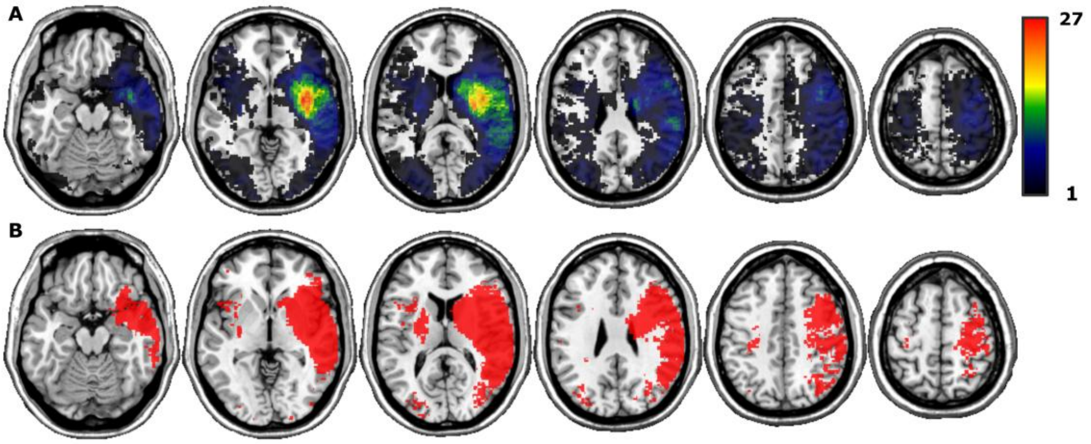
